# CDDO-Methyl Ester Inhibits BRAF Inhibitor Resistance and Remodels the Myeloid Compartment in BRAF-mutant Melanoma

**DOI:** 10.1101/2023.08.01.551524

**Authors:** Gretel M. Torres, Helen C. Jarnagin, Chanhyuk Park, Heetaek Yang, Noelle N. Kosarek, Rajan Bhandari, Chen-Yu Wang, Fred W. Kolling, Michael L. Whitfield, Mary Jo Turk, Karen T. Liby, Patricia A. Pioli

## Abstract

Approximately 50% of advanced melanomas harbor activating BRAF^V600E^ mutations that are sensitive to BRAF inhibition. However, the duration of the response to BRAF inhibitors (BRAFi) has been limited due to the development of acquired resistance, which is preceded by recruitment of immunosuppressive myeloid cells and regulatory T cells (T_regs_). While the addition of MAPK/ERK kinase 1 inhibitors (MEKi) prolongs therapeutic response to BRAF inhibition, most patients still develop resistance. Using a Braf^V600E/+^/Pten^-/-^ graft mouse model of melanoma, we now show that the addition of the methyl ester of the synthetic triterpenoid 2-cyano-3,12-dioxooleana-1,9(11)-dien-28-oic acid (C-Me) to the BRAFi vemurafenib analog PLX4720 at resistance significantly reduces tumor burden. Dual treatment remodels the BRAFi resistant-tumor microenvironment (TME), reducing infiltration of T_regs_ and tumor associated macrophages (TAMs), and attenuates immunosuppressive cytokine production. For the first time, we characterize myeloid populations using scRNA-seq in BRAFi-resistant tumors and demonstrate that restoration of therapeutic response is associated with significant changes in immune-activated myeloid subset representation. Collectively, these studies suggest that C-Me inhibits acquired resistance to BRAFi. Use of C-Me in combination with other therapies may both inhibit melanoma growth and enhance therapeutic responsiveness more broadly.

## INTRODUCTION

Although melanoma accounts for approximately 1% of all skin malignancies, it is responsible for the majority of skin cancer-related deaths *(1)*. Activating mutations in BRAF, a key serine-threonine kinase in the MAPK signaling pathway, occur in approximately 50% of cutaneous melanomas *(2)(3)*. While the use of targeted therapies such as vemurafenib to inhibit BRAF activation has led to significant advances in survival, many patients develop acquired resistance, typically within 6-8 months *(4)*. Although the first line use of combination BRAFi and MEKi therapy has led to significant improvements in survival, the development of secondary resistance and short duration of response (progression-free survival in only 19% of patients after 5 years) are persistent challenges in the effective treatment of melanoma patients *(5)*.

In addition to tumor cell-intrinsic factors, immune suppression in the tumor microenvironment (TME) is implicated in the regulation of therapeutic resistance. As mediators of inflammation, tumor-associated macrophages (TAMs) constitute a significant source of immune suppression in the TME, as they induce regulatory T cell (T_reg_) generation, programmed death-1 (PD-1)-dependent lymphocyte immunosuppression, and tumor-associated neo-angiogenesis (reviewed in *(6)*). Although short-term inhibitor treatment results in decreased CCL2 expression and loss of immunosuppressive myeloid cells from melanoma tumors, the BRAFi-resistant melanoma TME is characterized by restored baseline CCL2 expression, TAM and myeloid-derived suppressor cell (MDSC) repopulation, and reduced numbers of tumoral CD8 T cells *(7)*.

The triterpenoid 2-cyano-3,12-dioxooleana-1,9(11)-dien-28-oic acid methyl ester (C-Me) is a synthetic derivative of oleanoic acid (OA), which has anti-inflammatory and anti-carcinogenic properties *(8)*. C-Me has been shown to inhibit activation of signaling pathways aberrantly upregulated in cancer, including STAT3, MAPK, and PI3K *(9–11)*, and our prior work demonstrates that C-Me enhances TME immune activation by inhibiting myeloid cell tumor infiltration and redirecting TAM activation *(10, 12–14)*. In addition to remodeling the tumoral myeloid compartment, C-Me alters T cell distribution in tumors, increasing the ratio of CD8 to CD4 T cells and reducing T_reg_ infiltration in an estrogen receptor negative breast cancer model *(14)*.

Because the onset of BRAFi resistance is characterized by restoration of immune suppression and because C-Me attenuates tumoral immune suppression, we hypothesized that C-Me would inhibit BRAFi resistance and induce immune activation in the melanoma TME. To address this, we tested the therapeutic efficacy of monotherapy BRAFi vs. BRAFi/C-Me dual treatment. We now show that the addition of C-Me to the vemurafenib analog PLX4720 at the onset of resistance significantly reduces tumor burden and promotes immune activation. Tumors in mice that received combination therapy are characterized by significant reductions in TAM and T_reg_ infiltration. We also report for the first time that BRAFi-resistant tumors are populated by “Mrc1 Mac” and “CCR2 Mac” myeloid subsets, and that restoration of therapeutic response is associated with expansion of a “Cxcl2 Mac” myeloid population characterized by immune activation. Collectively, our results suggest that dual vemurafenib/C-Me treatment inhibits BRAFi resistance and converts immunologically “cold” TMEs into “hot” tumors.

## MATERIALS AND METHODS

### Mice

*Tyr::CreER*^*+/-*^*Braf*^CA/+^*Pten*^*lox/lox*^ (Braf/Pten) *(15)* were bred onto the C57BL/6 background. C57BL/6 mice were obtained from Charles River. Mice were handled ethically according to the Regulations for the Management of Laboratory Animals at the Geisel School of Medicine at Dartmouth. The experimental protocol for the ethical use of these animals was approved by the Institutional Animal Care and Use Committee at the Geisel School of Medicine at Dartmouth (protocol 00002010). Mice were euthanized by inhalation of carbon dioxide followed by cervical dislocation. All efforts were made to minimize animal suffering.

### Tumor Model and Treatment

C57BL/6 mice were dorsally grafted with ∼1cm^2^ tail skin section donated from Braf/Pten mice, and tumors were induced one week later by topical application of 4-hydroxy-tamoxifen (0.5ug/graft in DMSO)(Peprotech) at the graft site. Equal numbers of male and female age-matched mice were assigned to each cohort. Mice bearing palpable tumors (∼14 days post induction) were fed PLX4720 (BRAFi)-containing chow until the end of the experiment (Fig.1A). PLX4720 (MedChemExpress) was compounded in AIN-76A rodent diet (417 mg PLX4720/kg) by Research Diets, Inc. Tumors were deemed resistant to BRAFi after demonstrating sustained growth of >10% in skin thickness for 3 consecutive measurements. Resistant mice were then placed on mono-(BRAFi alone) or dual-therapy (BRAFi + C-Me; 50mg C-Me/kg) for 30 days and both tumors and dLNs were harvested. Investigators were not blinded during experimental monitoring, but blinding was applied during analysis of scRNA-seq, ELISA/multiplex, and immunoblot data.

### Flow Cytometry

Tumors were digested for 45 minutes with gentle shaking at 37°C in HBSS containing 7 mg/ml Collagenase D and 2 mg/ml DNase I (Sigma). Single-cell suspensions were stained with fluorophore-conjugated antibodies obtained from BioLegend: anti-CD206-Pe-Dazzle, anti-CD206-FITC, anti-CD11b-PerCP, anti-Gr-1-Pe/Cy7, anti-CD115-APC, anti-IA/IE-APC/Cy7, anti-FoxP3-FITC, anti-CD45-BV510, anti-CD45-APC/Cy7, anti-CD4-BV605, anti-CD4-PerCP, anti-CD8-BV711, anti-CD8-BV510, anti-CD3e-Pe/Cy5.5, anti-CD3e-BV421, anti-F4/80-BV421 and anti-FoxP3-Pe (eBioscience). Cells were stained for 1 hour at 4 °C with 2% rat serum (Sigma) to reduce antibody binding to Fc receptors. In all conditions, doublets and multiplets were excluded by forward scatter pulse width (SSC-W) vs. forward scatter pulse area (SSC-A) gating. Dead cells were eliminated by Fixable Blue Dead stain kit (Thermofisher Scientific) positivity. Gating of positively stained cells was determined by single-stain cell controls. Cells were analyzed using a 27-color ZE5 cell analyzer (Bio-Rad) with 5 laser sources (355nm, 488nm, 640nm, 405nm, 561nm) and FlowJo 9.8.1 (Treestar).

### Immunoblot

Nuclear lysates were prepared using NE-PER™ Nuclear and Cytoplasmic Extraction Reagents (Thermofisher Scientific). Lysates were analyzed for total protein concentration using a BCA assay kit (Thermofisher Scientific). Ten micrograms of each lysate were separated on Mini-PROTEAN TGS precast protein gels 4-15% (Bio-Rad) and electrotransferred to nitrocellulose membrane (GE Healthcare) in Tris-glycine buffer with 20% methanol. Membranes were blocked in 5% milk in 1xTBS and 0.05% Tween-20 (Santa Cruz) for 1hr at RT. Blots were probed with primary detection antibodies for s6 and p-s6(Cell Signaling). Primary antibodies for S6 and p-S6 were incubated at 1:1000 dilution in Tris-buffered saline with 0.1% Tween 20 and 5% w/v BSA overnight at 4°C with rotation. Following incubation with primary antibodies, blots were incubated with horseradish peroxidase-conjugated secondary antibodies (goat anti-mouse or goat anti-rabbit) (Bio-Rad) at a 1:2,000 dilution in Tris-buffered saline with Tween 20 (TBST) and 5% milk for 1hr at room temperature with gentle agitation on a rocker, followed by thorough washing in TBST. Blots were visualized using Pierce™ ECL Western Blotting Substrate (ThermoFisher Scientific).

### Enzyme-linked Immunosorbent Assay (ELISA) and Luminex

Secreted protein expression of tumor culture supernatants was quantified for CCL2, IL-6, IFN-γ, TNF-α, IL-10, IL-1α, and IL-1β by ELISA (Invitrogen). Protein levels of RANTES (CCL5), Eotaxin (CCL11), LIF, and GM-CSF were quantified by multiplex (Millipore) according to manufacturers’ protocols.

### Sample Preparation for Single Cell

Melanoma tumors were removed from male Braf/Pten mice six weeks post 4-hydroxy-tamoxifen topical application (0.5ug/mice in DMSO dorsally). Tumors were digested for 45 minutes with gentle shaking at 37°C in HBSS containing 3 mg/ml Collagenase IV and 2.8 mg/ml DNase-I (Sigma). Cells from BRAFi (n = 5) or BRAFi/C-Me (n = 6) tumors were pooled and CD45^+^ cells were isolated using CD45 MicroBeads (mouse, Miltenyi Biotec) according to the manufacturer’s protocol. Single cell suspensions were then tagged with mouse Total-Seq anti-mouse hashtags C0306 (Brafi: Hashtag 6) and C0307 (Brafi + C-Me: Hashtag 7) (BioLegend) according to manufacturers’ protocols.

### scRNA-seq and Bioinformatics

Concentration and viability of hashtagged single cell suspensions were measured using a Cellometer K2 instrument and AO/PI staining (Nexcelom Bioscience). An equal number of cells were mixed together prior to loading on a 10x Genomics Chromium NextGEM Chip G, targeting 10,000 cells for capture and processed to Illumina libraries following the 10x Genomics 3’ v3.1 protocol. Libraries were pooled and sequenced on an Illumina NextSeq2000 instrument to an average depth of 30,000 reads/cell.

Raw sequencing data were processed through Cell Ranger, version 6.0.1, from 10x Genomics. Downstream analysis was processed in R version 4.1.0 using the Seurat package version 4.0.5*(16)*. Cells were discarded if the unique feature counts were not between 200-2,500 and if they had more than 5% mitochondrial reads. Quality control filtering resulted in a total of 3495 single-cell transcriptomes.

These data were normalized and scaled using the NormalizeData(), FindVariableFeatures(), and ScaleData() functions. Principal component analysis dimensions were calculated with the RunPCA() function. Clustering of the data was performed with the FindNeighbors() and FindClusters() functions. Per interpretation of an Elbow Plot for this dataset, the first 10 PCA dimensions were used to run the FindNeighbors() function.

Cell clustering was accomplished by verifying the clustering significance through an Elbow Plot and by observing the clustering at different resolutions through the clustree package version 0.4.4*(17)*. We clustered the cells with the FindClusters() function using our optimized resolution of 0.25, as identified from our clustree plot. We then used the RunUMAP() function with previously set parameters and 10 PCA dimensions for visualization.

CITE-Seq cell hashing was used to separate the cells between treatment conditions. The reads for cell hashing were normalized in Seurat through the NormalizeData() function. We then identified which cluster had significant enrichment of the condition-designation hashes using the FindAllMarkers() function. We then subset the dataset identifying cells that expressed the specific hashtag greater than 0.5 as BRAFi treated, BRAFi + C-Me treated, or not hashed.

To identify gene enrichment in each cell cluster, we used the FindAllMarkers() function for the dataset. This performs a Wilcoxon Rank Sum test to identify enriched genes from each cluster. The top 20 genes in each cluster were used to determine cell identity when comparing markers found in the literature for typical myeloid cell types.

GSEA analysis for each cluster was completed using the fgsea package version 1.18.0*(18)*. For each cluster, a Wilcox Area Under the Curve analysis was conducted to calculate the expression enrichment within each cluster for all genes. These genes were then ranked by auc value and logFC value for each cluster. Using the fgsea() function, these ranked genes were compared to the pre-weighted genes within the Hallmark pathways of the GSEA database. The most significant core enrichment genes were identified by performing the FindAllMarkers() function on the prescribed set of core enrichment genes through GSEA.

### Statistical Analyses

Figures are representative of at least three independent experiments as indicated in Figure Legends. All experiments were repeated at least three times, unless otherwise noted. Biological and experimental replicates are indicated in Figure Legends (n). Results are described as mean ± SE and were analyzed by unpaired student’s t-Test, unless otherwise noted, using GraphPad Prism 8. Outliers were excluded after performing a Grubb’s test (ESD method) with α = 0.05. Significance was achieved at p < 0.05.

## RESULTS

### BRAFi/C-Me dual treatment at the onset of BRAFi-resistance inhibits tumor growth

Although short-term treatment with BRAFi results in decreased CCL2 expression and loss of immunosuppressive myeloid cells from melanoma tumors, long-term treatment with BRAFi results in the development of resistance, which is characterized by the restoration of baseline CCL2 expression and TAM repopulation *(7)*. Because C-Me inhibits myeloid cell migration and significantly attenuates CCL2 production *in vitro* and *in vivo (10, 13, 14)*, we hypothesized that C-Me would counter vemurafenib resistance through relief of TME immune suppression.

To test this hypothesis, C57BL/6 mice were dorsally grafted with ∼1cm^2^ tail skin donated from Braf/Pten mice. Tumors were induced one week later by topical application of 4-hydroxy-tamoxifen (0.5mg/graft in DMSO) at the graft site. Mice bearing palpable tumors (∼14 days post induction) were fed chow containing PLX4720. Tumor thickness was measured weekly, and mice were deemed resistant to BRAFi after demonstrating sustained growth (>10% increase in skin thickness for 3 consecutive measurements). Tumors and dLNs were harvested from C57BL/6 mice bearing Braf/Pten tumors treated with mono-(BRAFi) or dual-therapy (BRAFi + C-Me) for 30 days after the development of resistance (Fig. 1A). As shown in Figure 1B, mice that received dual therapy showed significant reductions in tumor weight compared with mice that received BRAFi alone.

**Figure 1:**
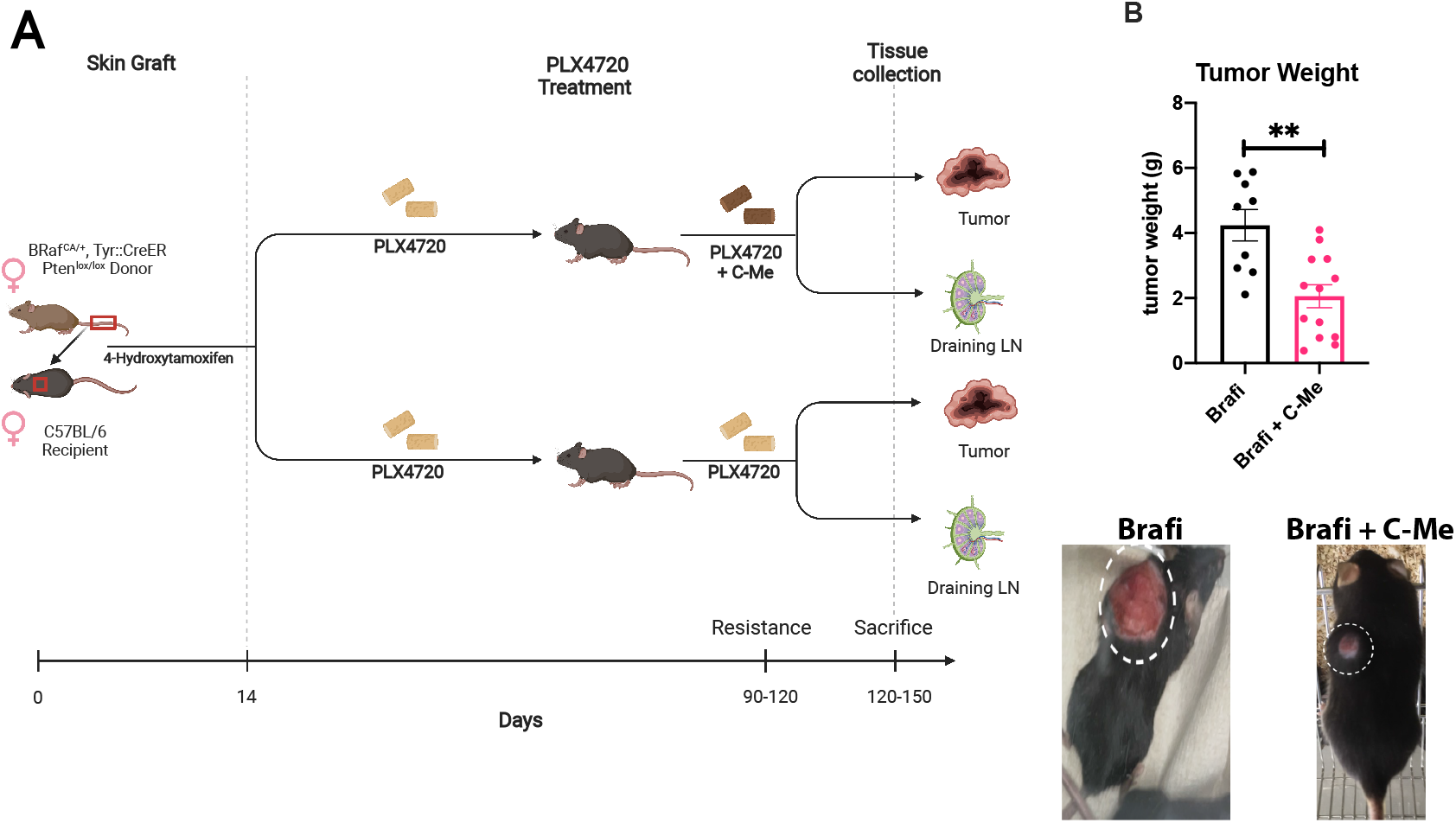
BRAFi/C-Me dual treatment after onset of BRAFi-resistance reduces tumor weight. (A)Tumor model; C57BL/6 mice were dorsally grafted with ∼1cm^2^ tail skin section donated from Braf/Pten mice, and tumors were induced one week later by topical application of 4-hydroxy-tamoxifen (0.5ug/graft in DMSO) at the graft site. Mice bearing palpable tumors (∼14 days post induction) were fed chow containing PLX4720 (417mg/kg). Tumors were deemed resistant to BRAFi after demonstrating sustained growth of >10% in skin thickness on 3 consecutive measurements. Resistant mice were then placed on a mono-(BRAFi) or dual-therapy (BRAFi + C-Me)(50mg C-Me/kg) for 30 days and both tumors and dLNs were harvested. (B) Tumor weights at 30 days post resistance and representative images. Tumor circled by dotted white lines. (A-B) Data were pooled from 3 independent experiments involving 3-5 female mice per group. No significant changes were observed on total body weight (not shown). Statistical significance was calculated by a two-tailed t-test. **p<0.01.

### Addition of C-Me to BRAFi post-resistance reduces numbers of T_regs_ and TAMs in tumors

Given the role imputed to myeloid and T cells in the development of resistance, we next interrogated how BRAFi/C-Me dual therapy altered immune cell representation in the dLN and tumors of treated mice. Since T_reg_ egress from the dLN during melanoma progression has been reported to promote immune suppression in Braf^V600E^ mutant tumors *(19)*, T cell subsets were analyzed using flow cytometry (Figs. 2A and 2B). As demonstrated in Figure 2C, CD4^+^ T cells account for the majority of the reduced tumoral T cell representation observed with dual therapy. Numbers of total CD3^+^ and CD8^+^ T cells as well as FoxP3^+^ T_regs_ (Fig. 2C) were significantly lower in dLNs from mice fed BRAFi/C-Me compared with BRAFi alone. The ratios of CD4^+^:CD8^+^ and CD4^+^ FoxP3^+^: CD8^+^ T cells were also significantly decreased in these dLNs (Fig. 2C), indicating that the representation of CD8^+^ T cells relative to CD4^+^ is enhanced with dual treatment.

**Figure 2:**
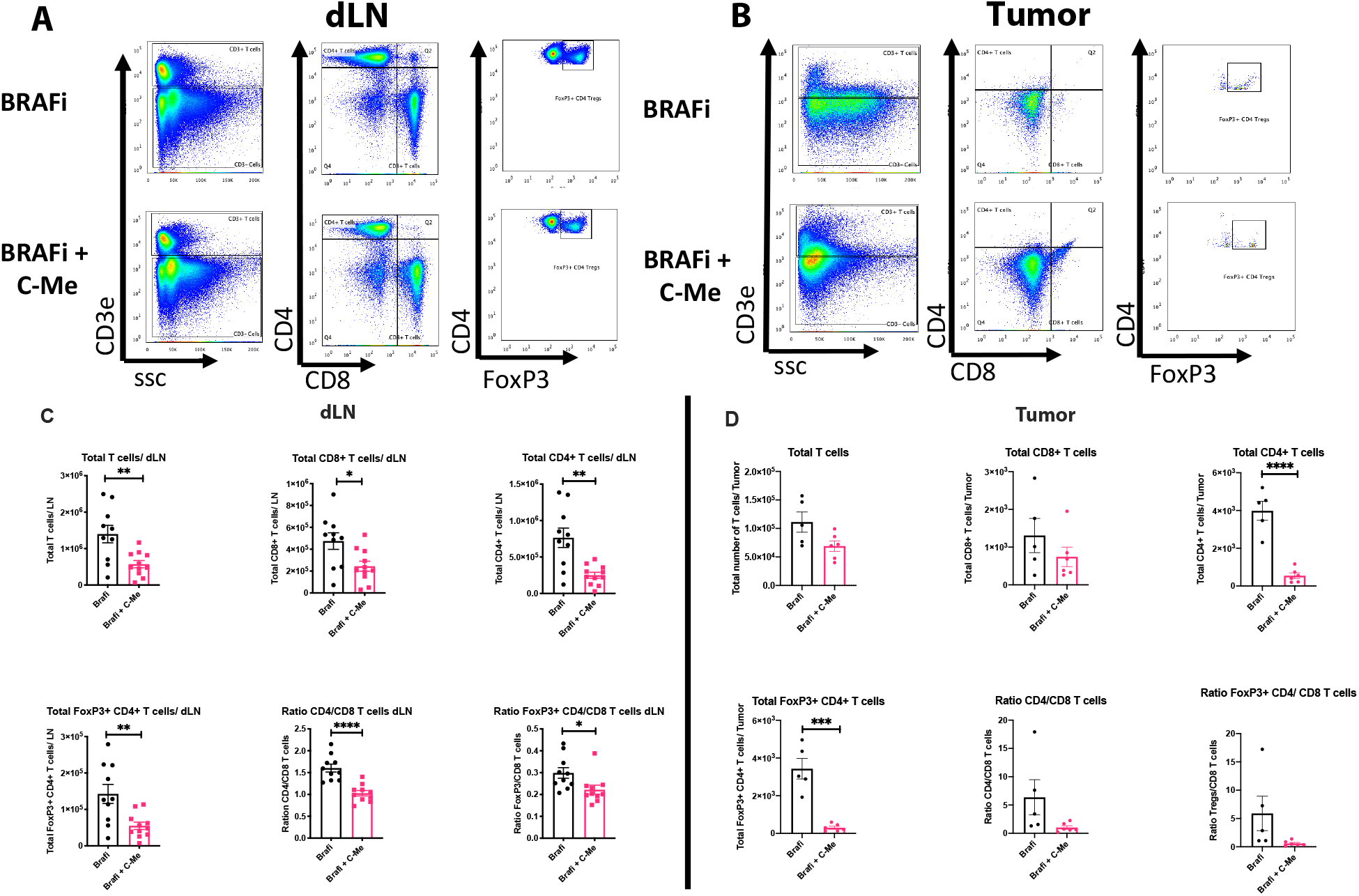
Dual treatment of BRAFi-resistant mice reduces T_regs_ in tumors and dLNs. (A-B) Gating strategy (doublets and dead cells removed prior). Numbers in plots represent percentages in each quadrant. (C) dLNs and (D) tumors from mono- (BRAFi) or dual-therapy (BRAFi + C-Me)(50mg C-Me/kg) treated mice were harvested and analyzed by multicolor flow cytometry to profile T cell populations. (C) Data were pooled from 3 independent experiments involving 5 mice per group. (D) Representative data from 3 independent experiments involving 5-6 mice per group. Statistical significance was calculated by two-tailed t-test. *p<0.05, **p<0.005, ****p<0.0001

Consistent with observations in the dLN, total CD4^+^ T cell and FoxP3^+^ T_reg_ numbers were significantly reduced in tumors of BRAFi/C-Me treated mice (Fig. 2D). The majority (∼95%) of the CD4^+^ T cells were also FoxP3^+^ T_regs_, regardless of treatment. These results suggest C-Me impedes the restoration of T_regs_ to BRAFi-resistant melanomas.

Because restoration of immunosuppressive myeloid cells is a hallmark of BRAFi resistance *(7)*, we next investigated the effect of BRAFi and C-Me on the macrophage compartment of the dLN and TME (Fig. 3A). TAMs were defined as IA/IE^+^, CD45^+^, CD11b^+^, F4/80^+^ cells (Fig. 3A). As observed in Figures 3B and 3C, infiltration of total CD206^+^ (MRC1) and CD115^+^ (M-CSFR) TAMs was significantly reduced in both tumors and dLNs of mice that received BRAFi and C-Me. In accordance with a prior report *(20)*, the proportion of MDSCs was not significantly altered by C-Me treatment (S.1 A-B).

**Figure 3:**
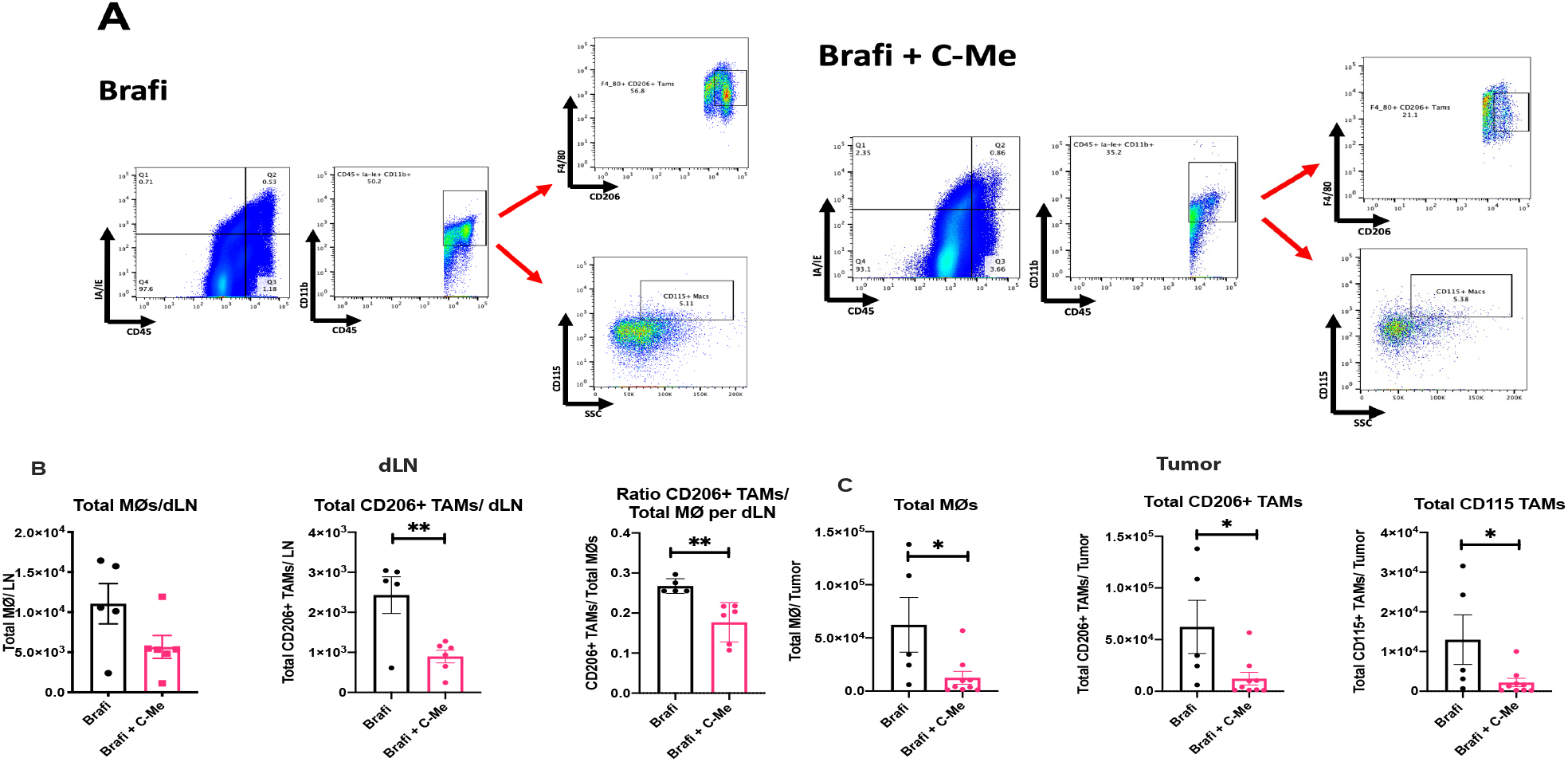
Treatment with BRAFi/C-Me at the onset of resistance reduces TAM numbers in tumors and dLNs. (A) Gating strategy (doublets and dead cells have been removed). Numbers in plots represent percentages in each quadrant. (B) dLNs and (C) tumors from mono- (BRAFi) or dual-therapy (BRAFi + C-Me)(50mg C-Me/kg) treated mice were harvested and analyzed by multicolor flow cytometry for examination of the TAM populations. (B) Data representative from 3 independent experiments involving 5-6 mice per group. (C) Data representative from 3 independent experiments involving 5-9 mice per group. Statistical significance was calculated by two-tailed t-test. *p<0.05, **p<0.005.

### C-Me induces immune activation and inhibits immune suppression in the TME of BRAFi-resistant mice

Because combination therapy inhibited immunosuppressive T_reg_ and TAM representation in the melanoma TME, we reasoned that these cellular changes would be reflected in altered cytokine production. To assess this, equal volumes (∼200 mgs) of tumor tissue were excised from mice that received mono or dual therapy and cultured *ex vivo* with complete RPMI in absence of drug. Supernatants were collected from tissue explants after 24, 48, and 96 hrs. Consistent with decreased TAM recruitment (Fig. 3), C-Me significantly attenuated tumoral CCL2 release (Fig. 4A, C-D) for at least 96 hrs *ex vivo*. Additionally, secretion of RANTES (CCL5) and eotaxin (CCL11), which are associated with endothelial cell chemotaxis and mediate epithelial-to-mesenchymal transition (EMT) *(21–23)*, was inhibited for at least 24 hrs *ex vivo* (Fig. 4A). As demonstrated in Figures 4B-D, levels of IL-6, LIF, VEGF, and IL-10, which are increased in human BRAF mutant melanomas and are associated with poor clinical outcome *(24)*, were also significantly decreased by dual BRAFi/C-Me treatment. Notably, expression of immune-activating IL-1α and IL-1β was induced by dual therapy *(25–30)* (Fig. 4 B-D), and although not statistically significant, increasing trends of TNF expression were also noted (Fig.4B). Collectively, these results suggest C-Me converts immunologically “cold” TMEs into “hot” states.

**Figure 4:**
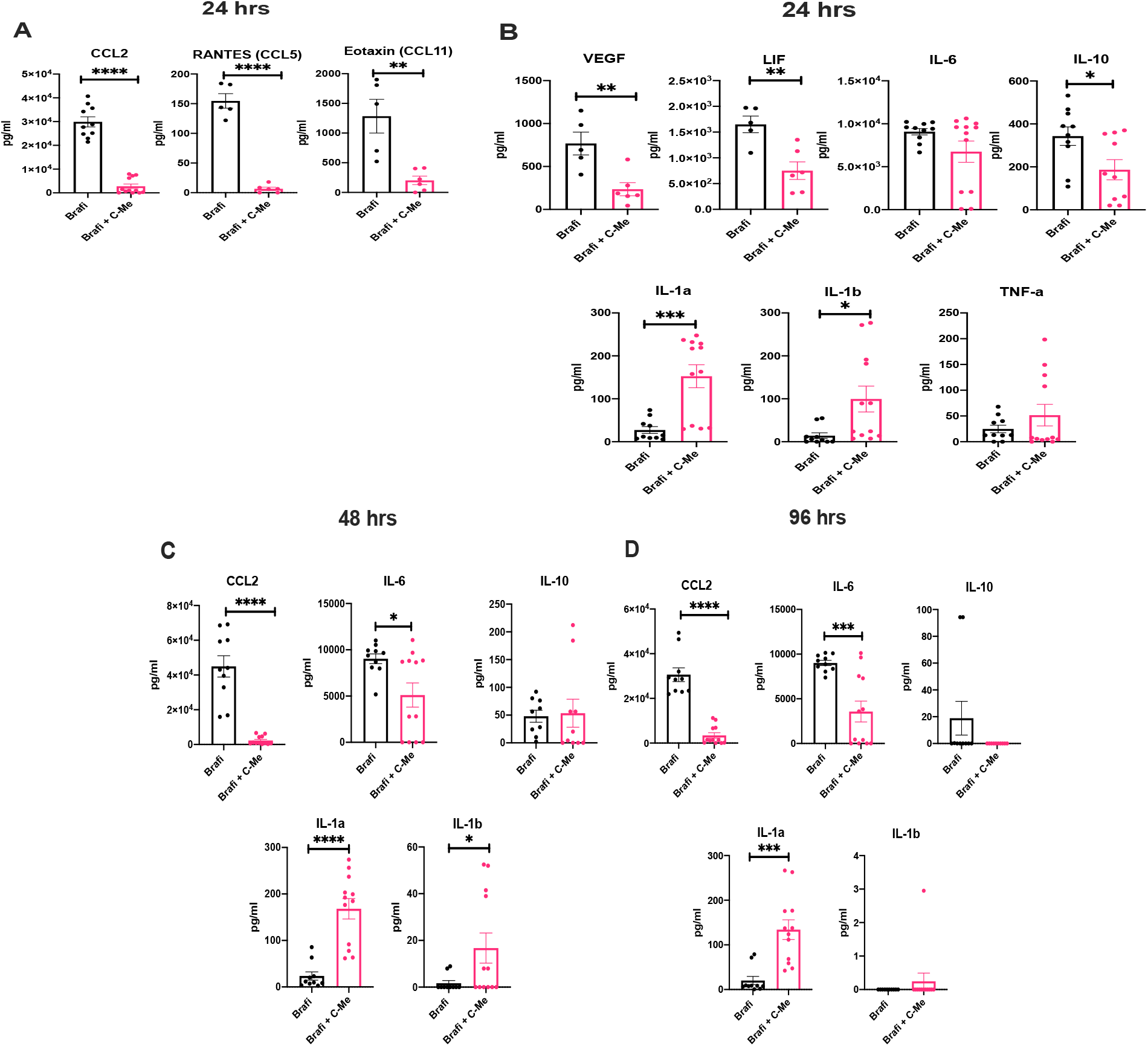
Addition of C-Me to BRAFi attenuates immunosuppressive cytokine and chemokine production in the TME, while inducing the production of immune-stimulatory cytokines up to 96 hrs *ex-vivo*. Tumors from mono- (BRAFi) or dual-therapy (BRAFi + C-Me)(50mg C-Me/kg) treated mice were sectioned (∼200mg) and cultured in complete RPMI. Supernatant from cultures tumors was collected at (A-B) 24 hrs, (C) 48 hrs, and (D) 96 hrs for Luminex or ELISA analysis cytokines and chemokines of interest. Representative from 3 independent experiments involving 5-6 mice per group. Statistical significance was calculated by two-tailed t-test. *p<0.05, **p<0.005, ***p<0.0005, ****p<0.0001.

### scRNA-seq identifies C-Me-induced changes in tumor immune subsets

Prior studies have documented TAM heterogeneity in the Braf/Pten model *(31)*, suggesting polyfunctionality among TAM subsets. Moreover, our results (Figs. 2, 3, and 4) indicated that the addition of C-Me to BRAFi at the onset of resistance alters the TME and T cell and TAM representation in the TME. To better define treatment effects on the tumor immune compartment at resistance, we performed scRNA-seq analysis (Fig. 5 A-F). Tumors from mice that received BRAFi alone or combination BRAFi/C-Me were enzymatically and mechanically dissociated into single cells, and CD45^+^ cells were isolated for transcriptional profiling using antibody-bound magnetic beads.

**Figure 5:**
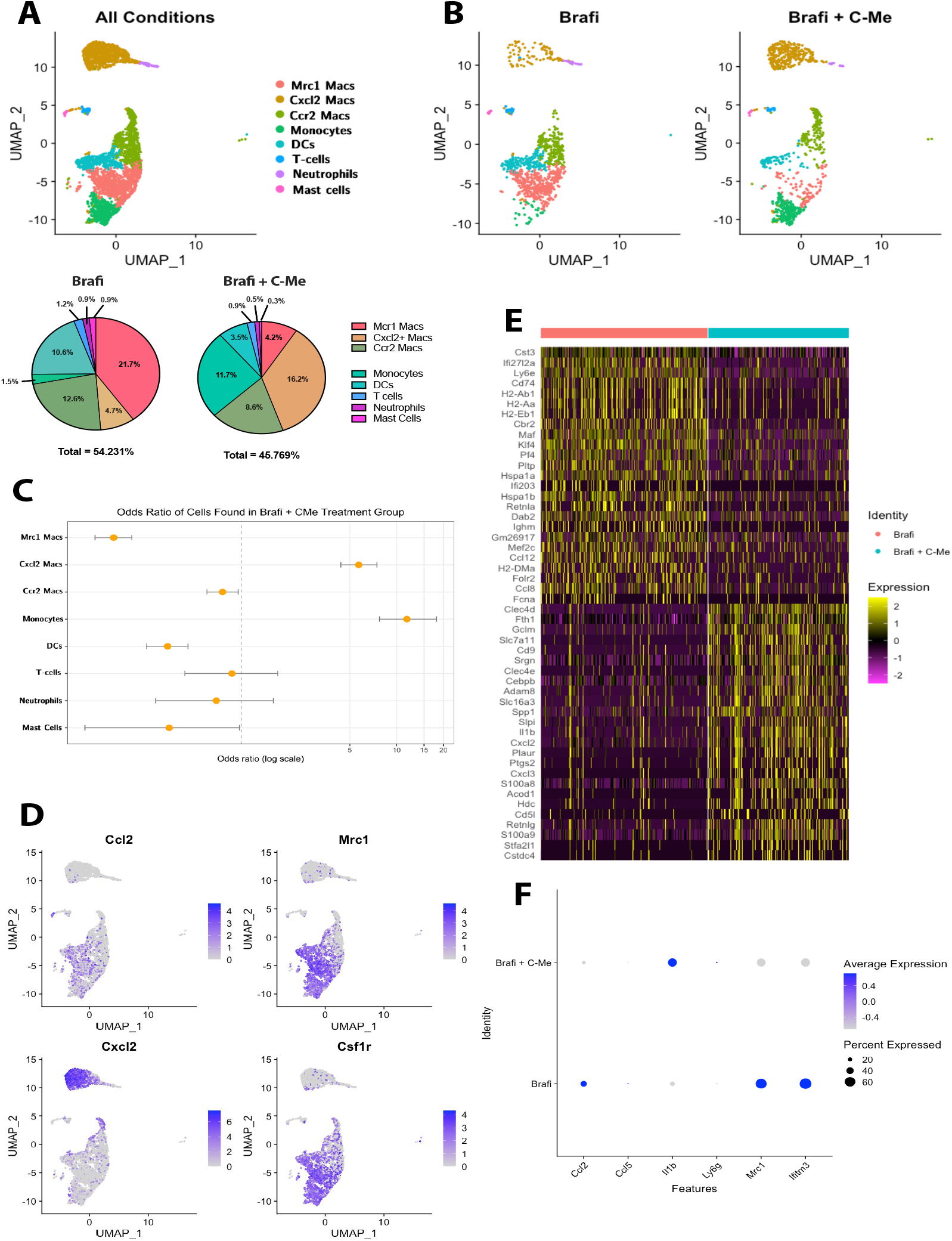
Global changes in myeloid cell populations of Brafi or Brafi/C-Me treated tumors. (A-B) Uniform manifold approximation and projection (UMAP) plots depicting single-cell transcriptomes of total (A) CD45+ cells and cells separated by (B) Brafi or Brafi/C-Me treatment (total cells n=3495). Proportional differences clustered cells between treatment conditions. (C) Proportional differences between cell types between treatment groups were confirmed with odds ratio testing. (D) Relative expression of *Ccl2, Mrc1, Cxcl2*, and *Csf1r* projected onto the UMAP plot to illustrate expression enrichment across cell populations. (E) Heatmap showing the 25 most differentially expressed genes across treatment conditions. See Table S2 for all differentially expressed genes between treatments. (F) Dot plot highlighting the average expression (color scale) and percentage of cells expressing (dot size) TAM-associated genes across treatment conditions.

Single-cell suspensions from tumors were used to construct scRNA-seq libraries following the 10X Genomics single-cell 3’RNA library construction and sequencing pipeline *(16–18)*. We sequenced 5,998 cells at a read depth of 31,936 reads per cell. Post-data processing resulted in 32,285 unique transcripts from 3,495 cells. To define tumoral immune populations, we computationally pooled data from CD45^+^ cells derived from tumors of 5 mice that received BRAFi monotherapy and 6 mice that received BRAFi/C-Me and used graph-based clustering to identify transcriptional subsets consisting of individual cell types. Comparison with the ImmGen database and established cell type markers identified four distinct monocyte/macrophage clusters and smaller clusters of dendritic cells (DCs), neutrophils, T cells, and mast cells. These findings are consistent with flow cytometry results, in which macrophages constituted the dominant immune cell population in tumors, representing approximately 70% and 40% of the total CD45^+^ cells in tumors with monotherapy and combination therapy, respectively (S.2).

Analysis of global changes in gene expression demonstrated that mice treated with BRAFi/C-Me were enriched for expression of *Il1b*, which has been shown to promote tumor regression by augmenting the production of tumor-experienced T cells, and *Cd9* and *Srgn*, which are associated with immune activation and indicative of favorable prognosis in cutaneous melanoma and other cancers *(28, 32, 33)*. In contrast, expression of *Ccl2, Mrc1, Ifitm3, Maf*, and *Klf4*, which have been implicated in the regulation of cancer growth, invasion, and metastasis (Fig. 5D-F), were higher in tumors from mice treated with BRAFi alone *(34–36)*.

### C-Me/BRAFi dual therapy at resistance remodels the myeloid compartment in tumors

To improve resolution and better define treatment-dependent differences in myeloid subsets in tumors, we computed percent differences in the monocyte/macrophage clusters based on treatment. Differential expression analysis was performed on significantly enriched/depleted populations to calculate cell type-specific gene expression changes between the two treatment arms.

As observed in Figures 5A-B and D, combined BRAFi/C-Me treatment was associated with enrichment of a myeloid subset characterized by expression of *Cxcl2* and *Il1b*; this subset was designated “Cxcl2 Mac” and is depicted in gold. An odds ratio test was used to confirm proportional differences in clusters based on treatment group, which demonstrated that “Cxcl2 Mac” and “Monocytes” were 5.71 and 11.7 times more likely to be found in BRAFi/C-Me vs. BRAFi alone treatment groups (CI 4.39-7.48 and CI 7.77-18.02). Conversely, cells in the “Mrc1 Mac” cluster (depicted in pink) were 6.59 times more likely to be associated with monotherapy compared with dual therapy (CI 5.05-8.7) (Fig. 5C). The “Mrc1 Mac” cluster was highly enriched for *Ccl2, Csf1r, and Mrc1*, which are commonly associated with immunosuppressive TAMs (Fig. 5D, F). DC, T cell, and mast cell clusters were not preferentially enriched based on treatment (Fig. 5A-C).

To define the transcriptional profiles of the myeloid subsets, we surveyed the top 10 differentially expressed genes in each cluster (Fig. 6A, Table S1). In addition to *Cxcl2* and *Il1b*, expression of *Srgn, Stfa2l1*, and *S100a8/a9*, which regulate inflammation, cytokine release, and antigen presentation was increased in the “Cxcl2 Mac” subset, which is enriched with BRAFi/C-Me treatment *(37, 38)* (Fig. 6A-B). The “Mrc1 Mac” cluster, which is expanded with BRAFi monotherapy, was characterized by high expression of *Maf*, a transcriptional regulator of immunosuppressive macrophage polarization that binds to a conserved noncoding region of the *csf-1r* gene *(34)*. Both Mrc1 and Maf are overexpressed in cancer and are associated with poor patient prognosis *(34, 39). Ccl8*, which recruits T_regs_ *(40)*, and *Ccl12*, which mediates monocyte recruitment, were also highly expressed by the “Mrc1 Mac” cluster (Fig. 6A-B) *(41)*.

**Figure 6:**
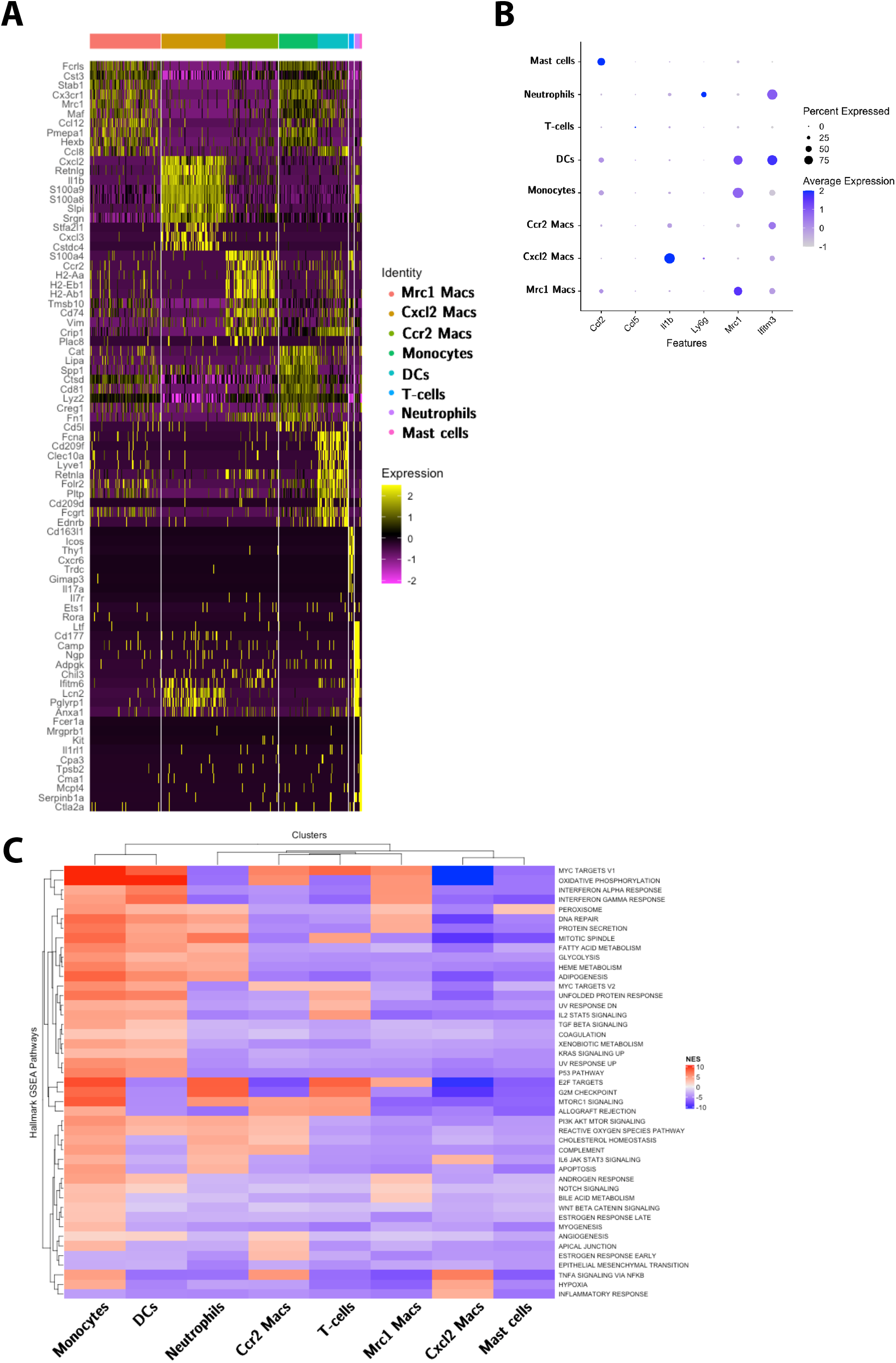
Cell type-specific expression changes induced by drug treatment. (A) Heatmap showing the 10 most differentially expressed genes within each cluster, as indicated by Seurat. All differentially expressed genes per cluster are shown in Table S1. (B) Dot-plot illustrating enrichment of key TAM-associated genes within each cell cluster. (C) Heatmap of statistically significant normalized enrichment scores (NES) for the GSEA hallmark pathways in each cell cluster across all treatment conditions (p-value<0.05). Dendrograms for Euclidean distance of NES for clusters and Hallmark Pathways identified.

Given the treatment-dependent changes observed in myeloid subset representation, gene set enrichment analysis (GSEA) was used to identify differences among the clusters in pathway regulation. As demonstrated in Figure 6C, GSEA revealed that TNF signaling and inflammatory response signatures were enriched in the “Cxcl2 Mac” subset compared with other clusters. In contrast, the “Mrc1 Mac” cluster displayed upregulation of pathways associated with oxidative phosphorylation (OXPHOS), the interferon-α response (IFN-α), and Myc targets.

### Core genes of hallmark pathways within “Mrc1 Mac” subset support tumor progression, metastasis, and invasion, while the “Cxcl2 Mac” cluster supports immune activation and tumor growth arrest

Because the IFN-α pathway has opposing tumoricidal and immunosuppressive functions *(42)*, we interrogated the IFN-a-responsive genes upregulated in the “Mrc1 Mac” cluster (Fig. 7A). We observed induction of *Lpar6, Lap3, Lgals3bp, Epsti1, B2m, Bst2*, previously demonstrated to mediate cancer progression, invasion, and metastasis (Fig. 7B) *(43–48)*, and consistent with a tumor-promoting phenotype of the “Mrc1 Mac” cluster *(43–48)*.

**Figure 7:**
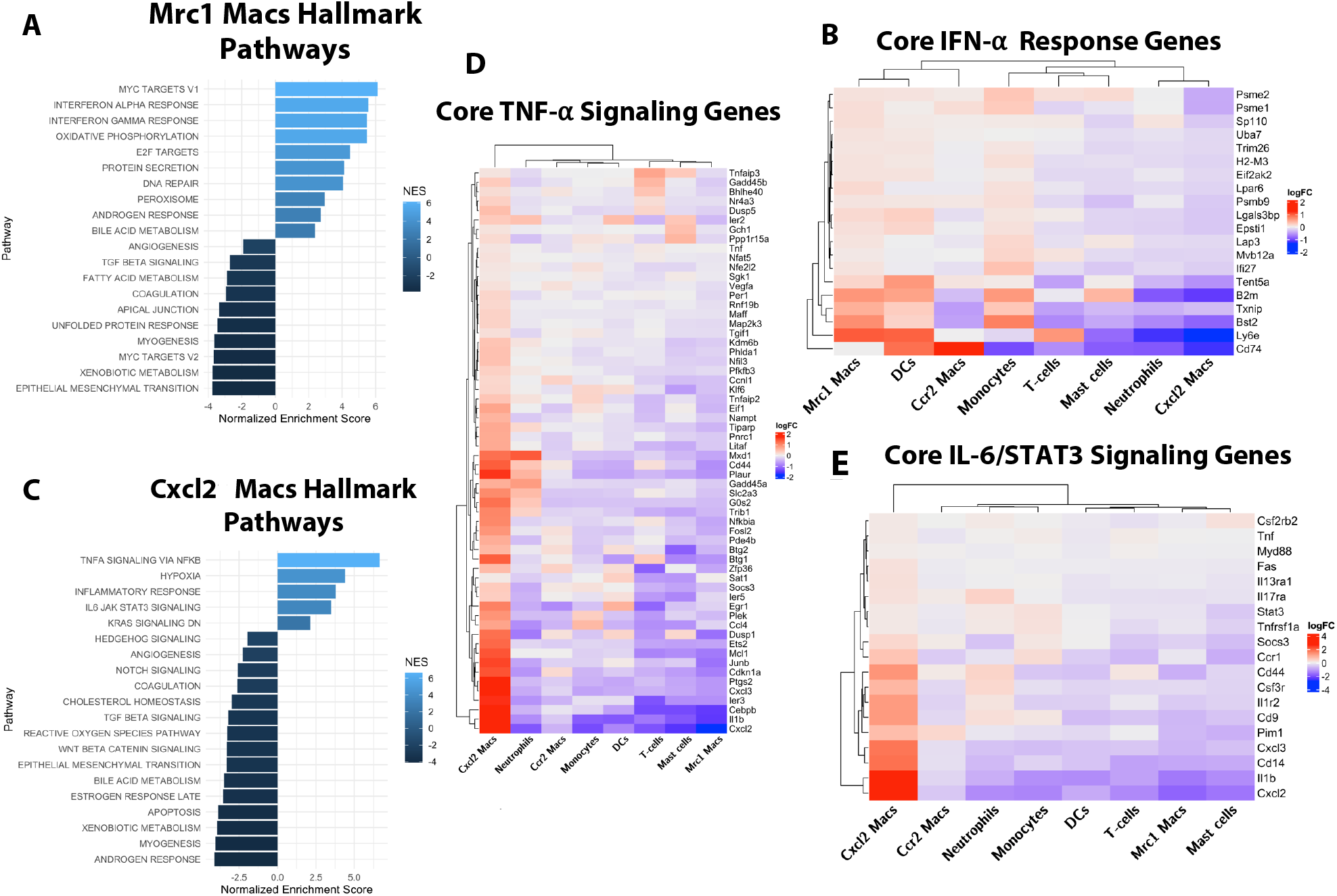
GSEA hallmark pathway analysis of “Mrc1 Mac” and “Cxcl2 Mac” clusters. (A, C) GSEA Normalized Enrichment Score plot for most statistically significant enriched pathways for (A) “Mrc1 Mac” and (C) “Cxcl2 Mac” (adjusted p-value <0.008). (B, D-E) LogFC of significant differentially-expressed core genes of the hallmark (B) INF-α response pathway enriched in the “Mrc1 Mac” cluster, (D)TNF Signaling via NFKB pathway, and (E) IL-6/JAK/STAT3 signaling pathway enriched in the “Cxcl2 Mac” cluster. Dendrograms depict Euclidean distance of logFC for clusters and core genes.

Conversely, hallmark pathways associated with TNF /NFkB and IL-6/JAK/STAT3 signaling were enriched in the “Cxcl2 Mac” cluster that was highly represented in mice that received dual therapy (Fig. 7C). Core genes that comprised the TNF signaling signature include *Il1b, Junb, Socs3*, and *Plaur*, which mediate M1-like macrophage activation *(49–51)*, and *Litaf*, a transcription factor involved in TNF transcription and inflammatory cytokine gene regulation (Fig. 7D)*(52)*. Because C-Me prevents phosphorylation of STAT3 *(10, 11)* and inhibits IL-6 (Fig. 4C-D)*(10)*, the upregulation of the IL-6/STAT3 signature was unexpected. However, genes highly expressed in this pathway are consistent with immune activation and include *Il1b, Il1r2, Cd14*, and *Cxcl2* (Fig. 7E). Taken together, these results suggest that BRAFi/C-Me treatment not only impairs myeloid cell migration into tumors but also changes the overall myeloid transcriptional landscape to promote immune activation.

### Transcriptional analysis implicates *c-Maf* as a regulator of “Mrc1 Mac” subset activation

Single-cell analyses demonstrated that the addition of C-Me to BRAFi at resistance shifted the myeloid milieu, resulting in preferential expansion of the “Cxcl2 Mac” cluster and contraction of the “Mrc1 Mac” cluster. These changes suggest distinct cellular differentiation pathways underlie myeloid subset representation. To investigate a potential mechanism by which this occurs, we analyzed transcription factors (TFs) within the CD45^+^ tumor population as a whole and by cell cluster (Fig. 8 A-B). Consistent with Figure 5E, global expression analysis showed BRAFi only-treated tumors increased expression of *Maf, Klf4, Fos*, and *Fosb*, while BRAFi/C-Me treatment increased expression of *Junb*, which is required for IL-1b expression and contributes to TNF production *(51)*(Fig. 8A).

**Figure 8:**
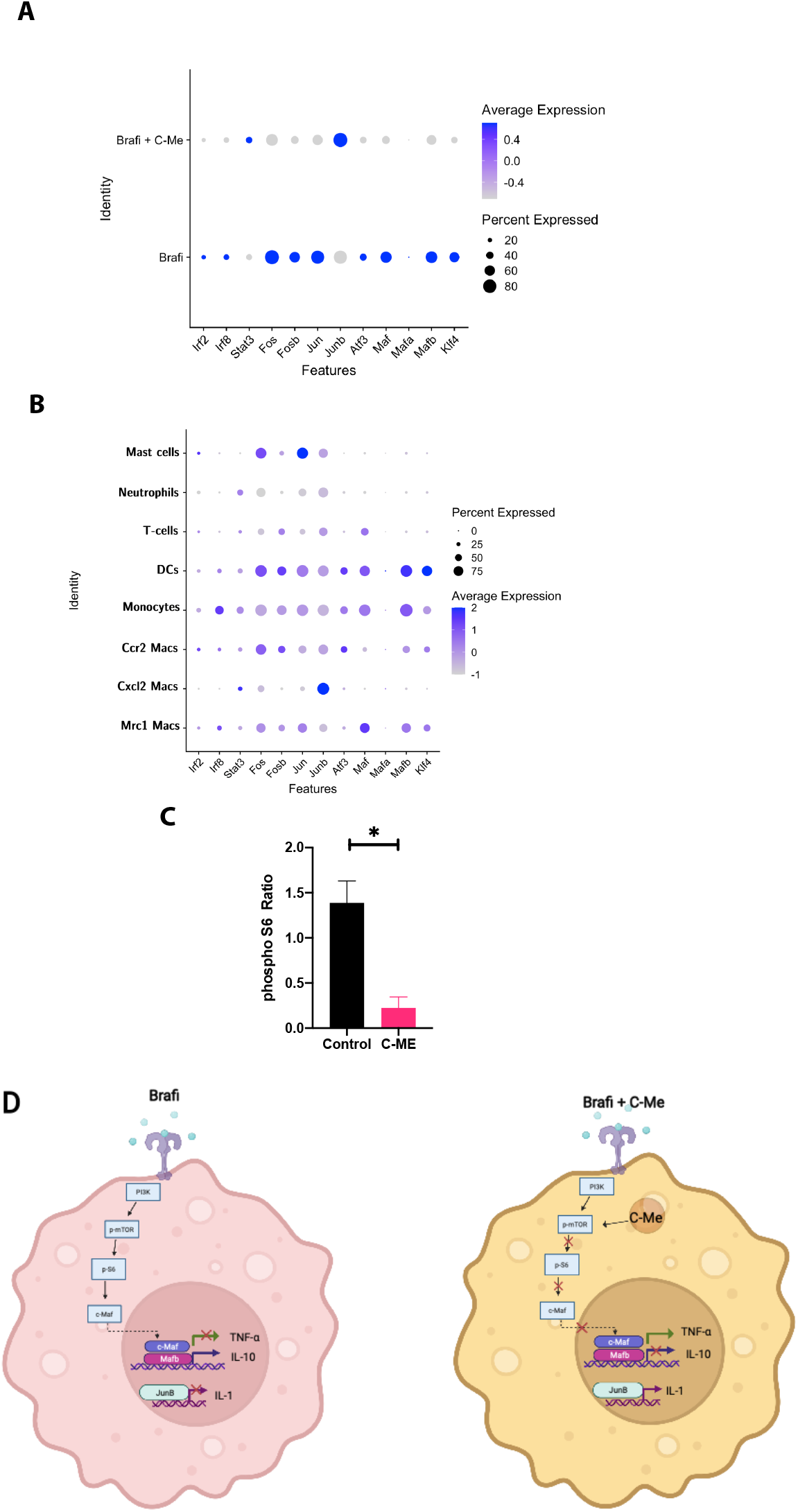
Differential expression of *MAF* and *JunB* in “Mrc1 Mac” and “Cxcl2 Mac” clusters. (A-B) Dot-plot illustrating enrichment of key transcription factors across (A) treatment conditions and (B) clusters. (C) Nuclear lysates were prepared from mouse BMDMs cultured for 5 days with tumor supernatants from *ex vivo* mono- or dual-therapy treated tumors. Lysates were quantified by BCA and 10mg total protein was loaded in each lane. Quantification of band intensity (ratio of pS6/S6) plotted. Representative from 3 independent experiments involving 3 tumors per group. Statistical significance was calculated by two-tailed t-test. *p<0.05 (D) Proposed model for C-Me mechanism of action in myeloid subset activation.

We next interrogated TF representation in the “Cxcl2 Mac” and “Mrc1 Mac” subsets, as cell numbers in these clusters shift dramatically dependent on drug treatment. As observed in Figure 8B, the “Mrc1 Mac” cluster contributed the majority of *Maf* expression, while the “Cxcl2 Mac” cluster was dominated by *JunB* (Fig. 8B). Thus, we hypothesized that C-Me inhibits mammalian target of rapamycin (mTOR) activation, resulting in decreased translation of Maf and IL-10. To address this, we investigated phosphorylation of the ribosomal protein S6, which is downstream of mTOR (Fig. 8D). Bone marrow-derived macrophages (BMDMs) were cultured for 5 days in conditioned media (CM) from tumors of mice treated *in vivo* with mono- or dual therapy. As shown in Figure 8C, S6 phosphorylation was significantly reduced in BMDMs cultured with CM from BRAFi/C-Me compared with BRAFi-only treated mice, supporting a role for C-Me in the regulation of mTOR activation.

## DISCUSSION

While previous studies have established tumor cell-intrinsic mechanisms of BRAFi resistance, myeloid heterogeneity and activation in the BRAFi-resistant melanoma TME are poorly understood. Using a well-characterized engrafted BRAF^V600E^/Pten^-/-^ melanoma mouse model, we have now shown that combining BRAFi with C-Me *in vivo* significantly attenuates hallmarks of vemurafenib resistance, including T_reg_ and TAM tumor infiltration and CCL2 production. Combination therapy at the onset of resistance reduced tumor weight, significantly inhibited tumor-promoting cytokine production, and remodeled the myeloid compartment in tumors. These studies highlight the potential of BRAFi/C-Me combination therapy to overcome BRAFi resistance and to promote immune activation in an immunologically “cold” TME.

Because C-Me was shown to induce disease stabilization in melanoma patients in a first-in-human phase I clinical trial *(53)*, our initial studies were performed using C-Me as a single therapeutic agent in inducible Braf/Pten mutant melanoma mouse models. However, the administration of C-Me as a monotherapy failed to arrest tumor growth or alter immune activation in the melanoma TME (S.3A-D). Intriguingly, we now show that although C-Me was ineffective during early tumor growth, the addition of C-Me to vemurafenib after the onset of resistance markedly attenuated tumor progression. This may be attributable to differences in the kinetics of tumor growth and immune cell representation in BRAFi-responsive vs. resistant TMEs. The molecular mechanism by which C-Me regulates gene expression is highly context-dependent *(54)*, and thus the clinical efficacy of this drug may be determined by local tissue microenvironments. Based on our results, we hypothesize this drug may be most efficient in combination with BRAFi at resistance onset and/or with other immunotherapies, including the use of immune checkpoint inhibitors.

Our studies were conducted with the Braf/Pten skin graft mouse model of melanoma, which recapitulates several features of human disease, including tumor stabilization for at least 3 months prior to resistance onset *(7)*. While the combination of BRAFi and C-Me significantly decreased tumor weight in mice bearing BRAFi-resistant tumors, tumor thickness did not differ dependent on treatment. Reductions in tumor circumference between mice treated with BRAFi/C-Me vs. BRAFi alone likely account for the difference in tumor weight. In addition to mediating changes in immune activation in the TME, C-Me has been shown to directly kill cancer cells by caspase-8 and TNF-dependent mechanisms *(55, 56)*. Thus, it is possible that observed changes in tumor growth may be attributable to increased tumor cell killing via enhanced TNF production as well as alterations in TAM and T cell activation and recruitment.

Recruitment of immunosuppressive immune cell subsets was inhibited by dual therapy. Notably, tumors of mice administered combination therapy had significantly fewer CD4^+^ T cells, which corresponded to a decrease in the T_reg_ population, with little change in total CD3^+^ and CD8^+^ T cell numbers. Similarly, tumor infiltration of CD206^+^ and CD115^+^ TAMs was also diminished. We hypothesize that these differences are attributable to C-Me-mediated changes in cytokine and chemokine production. In this regard, multiplex analysis demonstrated potent inhibition of secreted CCL2, CCL5, and IL-10. Consistent with our prior *in vitro* studies in a human melanoma culture model *(10)*, CCL2 production was nearly abrogated by the addition of C-Me. Significantly, the restoration of immunosuppressive myeloid cells at BRAFi resistance is contingent on MAPK reactivation and tumor cell production of CCL2 *(7)*. Thus, it is possible that one potential mechanism by which C-Me counters immunosuppressive myeloid cell recruitment and BRAFi resistance is through inhibition of tumoral CCL2 production. Moreover, reductions in IL-10 and CCL5 levels may contribute to impaired T_reg_ recruitment, as TAM-derived IL-10 contributes to tumoral T_reg_ accumulation, and CCL5 mediates T_reg_ recruitment *(57, 58)*.

While the production of VEGF, LIF, and IL-10 was significantly attenuated in conditioned media from dual-treated tumors, we also noted an increase in cytokines associated with immune activation, including IL-1α and IL-1β. These increases were sustained up to 96 hrs *ex vivo*, suggesting *in vivo* BRAFi/C-Me treatment may potentiate long-lived immune induction. Dependent on cell origin and local TME cytokine milieu, IL-1α and IL-1β can mediate either pro or anti-tumorigenic effects *(59)*. In combination with IL-2, IL-1β can promote differentiation of monocytes into “M1” macrophages and enhance antigen presentation *(59)*. Our prior work showed that C-Me increases IL-2 production by T cells, which may contribute to macrophage reprogramming *(10)*. Because the addition of C-Me inhibits tumor growth and remodels myeloid representation, we postulate that enhanced expression of IL-1 and TNF are indicative of enhanced immune activation in the previously “cold” BRAFi-resistant TME.

Notably, we observed a significant increase in monocyte representation in tumors treated with combination BRAFi/C-Me vs. BRAFi alone. Because we have shown that C-Me can re-educate TAMs *(10, 14)*, we hypothesize that these cells represent a transitional population, myeloid cells that are shifting from “Mrc1 Macs” toward “Cxcl2 Macs”. It is predicted that this precursor reservoir would become immune-activating within the TME. Alternatively, it is possible that C-Me recruits monocytes to tumors. However, given the dramatic reduction in CCL2 levels observed with the addition of C-Me, we predict that the most likely explanation for the observed increase in monocyte representation with dual treatment is redirection of myeloid activation. Ongoing studies in our laboratory are aimed at pursuing this answer.

Maf and MafB are critical regulators of immunosuppressive macrophage polarization, and they promote effector T cell suppression and enhance tumor progression in both mice and humans *(34, 60)*. Inhibition of c-Maf reduces IL-10 and VEGF levels and increases production of IL-1β and TNF. Furthermore, c-Maf KO macrophages show increased IFN-γ^+^ CD8^+^ T cell accumulation *(34)*. mTOR targets c-MAF translation and C-Me inhibits mTOR *(9, 61)*.

Transcription factor analysis showed tumors treated with BRAFi upregulated expression of c-Maf and MafB and Klf4, while expression of JunB was dominant with BRAFi/C-Me treatment, with relatively low levels of c-Maf/MafB expression. Notably, c-Maf expression in TAMs is associated with tumor growth and metastasis *(34)*, and expression of c-Maf is regulated by mTOR. Based on our results, we hypothesized that one potential mechanism by which combination BRAFi/C-Me reprograms myeloid activation in the melanoma BRAFi-resistant TME is through blockade of mTOR activation. In our proposed model (Fig. 8D), treatment with BRAFi/C-Me inhibits mTOR, resulting in decreased S6 phosphorylation, impaired translation of *c-Maf* and significant attenuation of immunosuppressive cytokine production. This leads to a shift in TF utilization, in which increased relative expression of *JunB* results in redirection of myeloid activation, relief of immune suppression, and the induction of immune activation in the BRAFi-resistant TME.

Collectively, our data suggest that combining vemurafenib with C-Me may inhibit acquired resistance in BRAF^v600E^ mutant melanoma. Because we have shown that C-Me induces immune activation in “cold” tumors, C-Me may have clinical utility as an adjuvant to enhance the efficacy of additional chemo-and immunotherapies in previously unresponsive tumors.

## ACKNOWLEDGEMENTS

This work was conducted with the Genomics and Molecular Biology Shared Resource (GMBSR) at Dartmouth, which is supported by NCI Cancer Center Support Grant 5P30CA023108 and NIH S10 (1S10OD030242) awards. Single cell studies were conducted through the Dartmouth Center for Quantitative Biology in collaboration with the GMBSR with support from NIGMS (P20GM130454) and NIH S10 (S10OD025235) awards. Flow cytometry and multiplex analysis were carried out in DartLab, the Immune Monitoring and Flow Cytometry Resource at the Norris Cotton Cancer Center at Dartmouth with NCI Cancer Center Support Grant 5P30CA023108-37.

## FUNDING

National Institutes of Health P20GM130454 (MLW, PAP)

National Institutes of Health R01CA226690 (KTL)

National Institutes of Health 5T32A1007363-28 (GMT)

Prouty Pilot Developmental Funds (PAP, MLW, MJT)

John Osborn Polak Endowment (RB)

## AUTHOR CONTRIBUTIONS

Conceptualization: GMT, MJT, PAP

Methodology: HCJ, NNK, FK IV

Investigation: GMT, HCJ, CP, HY, C-YW, RB

Funding Acquisition: MLW, KTL, PAP

Project Administration: PAP

Supervision: MLW, FK IV, MJT, KTL

Writing—original draft: GMT, HCJ, PAP

Writing—review and editing: GMT, HCJ, CP, HY, NNK, C-YW, RB, MLW, MJT, KTL, PAP

## DATA AND MATERIALS AVAILABILITY

Mouse single-cell RNA-seq data will be deposited at GEO and will be made publicly available. Flow cytometry data will be made available in Mendeley Data. This paper does not report original code. Any additional information required to reanalyze the data reported in this paper may be obtained from the lead contact upon request.

## REFERENCES

1. S. J. Henley, E. M. Ward, S. Scott, J. Ma, R. N. Anderson, A. U. Firth, C. C. Thomas, F. Islami, H. K. Weir, D. R. Lewis, R. L. Sherman, M. Wu, V. B. Benard, L. C. Richardson, A. Jemal, K. Cronin, B. A. Kohler, Annual report to the nation on the status of cancer, part I: National cancer statistics. Cancer 126, 2225–2249 (2020).

2. H. Davies, G. R. Bignell, C. Cox, P. Stephens, S. Edkins, S. Clegg, J. Teague, H. Woffendin, M. J. Garnett, W. Bottomley, N. Davis, E. Dicks, R. Ewing, Y. Floyd, K. Gray, S. Hall, R. Hawes, J. Hughes, V. Kosmidou, A. Menzies, C. Mould, A. Parker, C. Stevens, S. Watt, S. Hooper, R. Wilson, H. Jayatilake, B. A. Gusterson, C. Cooper, J. Shipley, D. Hargrave, K. Pritchard-Jones, N. Maitland, G. Chenevix-Trench, G. J. Riggins, D. D. Bigner, G. Palmieri, A. Cossu, A. Flanagan, A. Nicholson, J. W. C. Ho, S. Y. Leung, S. T. Yuen, B. L. Weber, H. F. Seigler, T. L. Darrow, H. Paterson, R. Marais, C. J. Marshall, R. Wooster, M. R. Stratton, P. A. Futreal, Mutations of the BRAF gene in human cancer. Nature 417, 949–954 (2002).

3. A. M. Menzies, L. E. Haydu, L. Visintin, M. S. Carlino, J. R. Howle, J. F. Thompson, R. F. Kefford, R. A. Scolyer, G. V. Long, Distinguishing Clinicopathologic Features of Patients with V600E and V600K BRAF - Mutant Metastatic Melanoma. Clin Cancer Res 18, 3242–3249 (2012).

4. J. Larkin, P. A. Ascierto, B. Dréno, V. Atkinson, G. Liszkay, M. Maio, M. Mandalà, L. Demidov, D. Stroyakovskiy, L. Thomas, L. de la Cruz-Merino, C. Dutriaux, C. Garbe, M. A. Sovak, I. Chang, N. Choong, S. P. Hack, G. A. McArthur, A. Ribas, Combined Vemurafenib and Cobimetinib in BRAF -Mutated Melanoma. N Engl J Med 371, 1867–1876 (2014).

5. C. Robert, J. J. Grob, D. Stroyakovskiy, B. Karaszewska, A. Hauschild, E. Levchenko, V. Chiarion Sileni, J. Schachter, C. Garbe, I. Bondarenko, H. Gogas, M. Mandalá, J. B. A. G. Haanen, C. Lebbé, A. Mackiewicz, P. Rutkowski, P. D. Nathan, A. Ribas, M. A. Davies, K. T. Flaherty, P. Burgess, M. Tan, E. Gasal, M. Voi, D. Schadendorf, G. V. Long, Five-Year Outcomes with Dabrafenib plus Trametinib in Metastatic Melanoma. N Engl J Med 381, 626–636 (2019).

6. M. Bied, W. W. Ho, F. Ginhoux, C. Blériot, Roles of macrophages in tumor development: a spatiotemporal perspective. Cell Mol Immunol (2023), doi:10.1038/s41423-023-01061-6.

7. S. M. Steinberg, T. B. Shabaneh, P. Zhang, V. Martyanov, Z. Li, B. T. Malik, T. A. Wood, A. Boni, A. Molodtsov, C. V. Angeles, T. J. Curiel, M. L. Whitfield, M. J. Turk, Myeloid Cells That Impair Immunotherapy Are Restored in Melanomas with Acquired Resistance to BRAF Inhibitors. Cancer Res 77, 1599–1610 (2017).

8. K. T. Liby, M. B. Sporn, T. A. Esbenshade, Ed. Synthetic Oleanane Triterpenoids: Multifunctional Drugs with a Broad Range of Applications for Prevention and Treatment of Chronic Disease. Pharmacol Rev 64, 972–1003 (2012).

9. X.-Y. Wang, X.-H. Zhang, L. Peng, Z. Liu, Y.-X. Yang, Z.-X. He, H.-W. Dang, S.-F. Zhou, Bardoxolone methyl (CDDO-Me or RTA402) induces cell cycle arrest, apoptosis and autophagy via PI3K/Akt/mTOR and p38 MAPK/Erk1/2 signaling pathways in K562 cells. Am J Transl Res 9, 4652–4672 (2017).

10. G. M. Torres, H. Yang, C. Park, P. A. Spezza, N. Khatwani, R. Bhandari, K. T. Liby, P. A. Pioli, T Cells and CDDO-Me Attenuate Immunosuppressive Activation of Human Melanoma-Conditioned Macrophages. Front. Immunol. 13, 768753 (2022).

11. Z. Duan, R. Y. Ames, M. Ryan, F. J. Hornicek, H. Mankin, M. V. Seiden, CDDO-Me, a synthetic triterpenoid, inhibits expression of IL-6 and Stat3 phosphorylation in multi-drug resistant ovarian cancer cells. Cancer Chemother Pharmacol 63, 681–689 (2009).

12. K. Tran, R. Risingsong, D. Royce, C. R. Williams, M. B. Sporn, K. Liby, The synthetic triterpenoid CDDO-methyl ester delays estrogen receptor-negative mammary carcinogenesis in polyoma middle T mice. Cancer Prev Res (Phila) 5, 726–734 (2012).

13. M. S. Ball, E. P. Shipman, H. Kim, K. T. Liby, P. A. Pioli, CDDO-Me Redirects Activation of Breast Tumor Associated Macrophages. PLoS One 11, e0149600 (2016).

14. M. S. Ball, CDDO-Me Alters the Tumor Microenvironment in Estrogen Receptor Negative Breast Cancer. Sci. Rep 10, 6560 (2020).

15. D. Dankort, D. P. Curley, R. A. Cartlidge, B. Nelson, A. N. Karnezis, W. E. Damsky, M. J. You, R. A. DePinho, M. McMahon, M. Bosenberg, Braf(V600E) cooperates with Pten loss to induce metastatic melanoma. Nat Genet 41, 544–552 (2009).

16. Y. Hao, S. Hao, E. Andersen-Nissen, W. M. Mauck, S. Zheng, A. Butler, M. J. Lee, A. J. Wilk, C. Darby, M. Zager, P. Hoffman, M. Stoeckius, E. Papalexi, E. P. Mimitou, J. Jain, A. Srivastava, T. Stuart, L. M. Fleming, B. Yeung, A. J. Rogers, J. M. McElrath, C. A. Blish, R. Gottardo, P. Smibert, R. Satija, Integrated analysis of multimodal single-cell data. Cell 184, 3573–3587.e29 (2021).

17. L. Zappia, A. Oshlack, Clustering trees: a visualization for evaluating clusterings at multiple resolutions. GigaScience 7, 1–9 (2018).

18. G. Korotkevich, V. Sukhov, A. Sergushichev, Fast gene set enrichment analysis. bioRxiv, 060012–060012 (2019).

19. T. B. Shabaneh, A. K. Molodtsov, S. M. Steinberg, P. Zhang, G. M. Torres, G. A. Mohamed, A. Boni, T. J. Curiel, C. V. Angeles, M. J. Turk, Oncogenic BRAF ^V600E^ Governs Regulatory T-cell Recruitment during Melanoma Tumorigenesis. Cancer Research 78, 5038–5049 (2018).

20. S. Nagaraj, J.-I. Youn, H. Weber, C. Iclozan, L. Lu, M. J. Cotter, C. Meyer, C. R. Becerra, M. Fishman, S. Antonia, M. B. Sporn, K. T. Liby, B. Rawal, J.-H. Lee, D. I. Gabrilovich, Anti-inflammatory Triterpenoid Blocks Immune Suppressive Function of MDSCs and Improves Immune Response in Cancer. Clin Cancer Res 16, 1812–1823 (2010).

21. R. Salcedo, H. A. Young, M. L. Ponce, J. M. Ward, H. K. Kleinman, W. J. Murphy, J. J. Oppenheim, Eotaxin (CCL11) Induces In Vivo Angiogenic Responses by Human CCR3 + Endothelial Cells. J Immunol 166, 7571–7578 (2001).

22. F. Zhu, P. Liu, J. Li, Y. Zhang, Eotaxin-1 promotes prostate cancer cell invasion via activation of the CCR3-ERK pathway and upregulation of MMP-3 expression. Oncology Reports 31, 2049–2054 (2014).

23. J. Ma, F. Shayiti, J. Ma, M. Wei, T. Hua, R. Zhang, J. Su, P. Chen, Tumor-associated macrophage-derived CCL5 promotes chemotherapy resistance and metastasis in prostatic cancer. Cell Biol Int 45, 2054–2062 (2021).

24. H. Sumimoto, F. Imabayashi, T. Iwata, Y. Kawakami, The BRAF–MAPK signaling pathway is essential for cancer-immune evasion in human melanoma cells. Journal of Experimental Medicine 203, 1651–1656 (2006).

25. M. K. Sgagias, A. Kasid, D. N. Danforth, Interleukin-1α and Tumor Necrosis Factor-α (TNFα) Inhibit Growth and Induce TNF Messenger RNA in MCF-7 Human Breast Cancer Cells. Molecular Endocrinology 5, 1740–1747 (1991).

26. S. L. Maund, W. W. Barclay, L. D. Hover, L. S. Axanova, G. Sui, J. D. Hipp, J. C. Fleet, A. Thorburn, S. D. Cramer, Interleukin-1α Mediates the Antiproliferative Effects of 1,25-Dihydroxyvitamin D _3_in Prostate Progenitor/Stem Cells. Cancer Res 71, 5276–5286 (2011).

27. S. L. Maund, L. Shi, S. D. Cramer, A. T. Slominski, Ed. A Role for Interleukin-1 Alpha in the 1,25 Dihydroxyvitamin D3 Response in Mammary Epithelial Cells. PLoS ONE 8, e81367 (2013).

28. R. J. North, R. H. Neubauer, J. J. Huang, R. C. Newton, S. E. Loveless, Interleukin 1-induced, T cell-mediated regression of immunogenic murine tumors. Requirement for an adequate level of already acquired host concomitant immunity. Journal of Experimental Medicine 168, 2031–2043 (1988).

29. O. A. W. Haabeth, K. B. Lorvik, H. Yagita, B. Bogen, A. Corthay, Interleukin-1 is required for cancer eradication mediated by tumor-specific Th1 cells. OncoImmunology 5, e1039763 (2016).

30. O. A. W. Haabeth, K. B. Lorvik, C. Hammarström, I. M. Donaldson, G. Haraldsen, B. Bogen, A. Corthay, Inflammation driven by tumour-specific Th1 cells protects against B-cell cancer. Nat Commun 2, 240 (2011).

31. C. J. Perry, A. R. Muñoz-Rojas, K. M. Meeth, L. N. Kellman, R. A. Amezquita, D. Thakral, V. Y. Du, J. X. Wang, W. Damsky, A. L. Kuhlmann, J. W. Sher, M. Bosenberg, K. Miller-Jensen, S. M. Kaech, Myeloid-targeted immunotherapies act in synergy to induce inflammation and antitumor immunity. Journal of Experimental Medicine 215, 877–893 (2018).

32. H. M. Koh, B. G. Jang, D. H. Lee, C. L. Hyun, Increased CD9 expression predicts favorable prognosis in human cancers: a systematic review and meta-analysis. Cancer Cell Int 21, 472 (2021).

33. X. Wang, H. Xiong, D. Liang, Z. Chen, X. Li, K. Zhang, The role of SRGN in the survival and immune infiltrates of skin cutaneous melanoma (SKCM) and SKCM-metastasis patients. BMC Cancer 20, 378 (2020).

34. M. Liu, Z. Tong, C. Ding, F. Luo, S. Wu, C. Wu, S. Albeituni, L. He, X. Hu, D. Tieri, E. C. Rouchka, M. Hamada, S. Takahashi, A. A. Gibb, G. Kloecker, H. Zhang, M. Bousamra, B. G. Hill, X. Zhang, J. Yan, Transcription factor c-Maf is a checkpoint that programs macrophages in lung cancer. Journal of Clinical Investigation 130, 2081–2096 (2020).

35. U. S. Rajapaksa, C. Jin, T. Dong, Malignancy and IFITM3: Friend or Foe? Front. Oncol. 10, 593245 (2020).

36. F. Yu, J. Li, H. Chen, J. Fu, S. Ray, S. Huang, H. Zheng, W. Ai, Kruppel-like factor 4 (KLF4) is required for maintenance of breast cancer stem cells and for cell migration and invasion. Oncogene 30, 2161–2172 (2011).

37. S. Wang, R. Song, Z. Wang, Z. Jing, S. Wang, J. Ma, S100A8/A9 in Inflammation. Front. Immunol. 9, 1298 (2018).

38. C.-S. Hsieh, P. deRoos, K. Honey, C. Beers, A. Y. Rudensky, A Role for Cathepsin L and Cathepsin S in Peptide Generation for MHC Class II Presentation. J Immunol 168, 2618–2625 (2002).

39. J. M. Debacker, O. Gondry, T. Lahoutte, M. Keyaerts, W. Huvenne, The Prognostic Value of CD206 in Solid Malignancies: A Systematic Review and Meta-Analysis. Cancers 13, 3422 (2021).

40. E. C. Halvorsen, M. J. Hamilton, A. Young, B. J. Wadsworth, N. E. LePard, H. N. Lee, N. Firmino, J. L. Collier, K. L. Bennewith, Maraviroc decreases CCL8-mediated migration of CCR5 + regulatory T cells and reduces metastatic tumor growth in the lungs. OncoImmunology 5, e1150398 (2016).

41. D. Argyle, T. Kitamura, Targeting Macrophage-Recruiting Chemokines as a Novel Therapeutic Strategy to Prevent the Progression of Solid Tumors. Front. Immunol. 9, 2629 (2018).

42. B. D.-M. Chen, F. Najor, Macrophage activation by interferon α + β is associated with a loss of proliferative capacity: Role of interferon α + β in the regulation of macrophage proliferation and function. Cellular Immunology 106, 343–354 (1987).

43. C. Fang, J. Zhang, H. Yang, L. Peng, K. Wang, Y. Wang, X. Zhao, H. Liu, C. Dou, L. Shi, C. Zhao, S. Liang, D. Li, X. Wang, Leucine aminopeptidase 3 promotes migration and invasion of breast cancer cells through upregulation of fascin and matrix metalloproteinases-2/9 expression. J Cell Biochem 120, 3611–3620 (2019).

44. X. Zheng, Y. Jia, L. Qiu, X. Zeng, L. Xu, M. Wei, C. Huang, C. Liu, L. Chen, J. Han, A potential target for liver cancer management, lysophosphatidic acid receptor 6 (LPAR6), is transcriptionally up-regulated by the NCOA3 coactivator. Journal of Biological Chemistry 295, 1474–1488 (2020).

45. E. Capone, S. Iacobelli, G. Sala, Role of galectin 3 binding protein in cancer progression: a potential novel therapeutic target. J Transl Med 19, 405 (2021).

46. T. Li, H. Lu, C. Shen, S. K. Lahiri, M. S. Wason, D. Mukherjee, L. Yu, J. Zhao, Identification of epithelial stromal interaction 1 as a novel effector downstream of Krüppel-like factor 8 in breast cancer invasion and metastasis. Oncogene 33, 4746–4755 (2014).

47. H. Zhang, B. Cui, Y. Zhou, X. Wang, W. Wu, Z. Wang, Z. Dai, Q. Cheng, K. Yang, B2M overexpression correlates with malignancy and immune signatures in human gliomas. Sci Rep 11, 5045 (2021).

48. W. D. Mahauad-Fernandez, W. Naushad, T. D. Panzner, A. Bashir, G. Lal, C. M. Okeoma, BST-2 promotes survival in circulation and pulmonary metastatic seeding of breast cancer cells. Sci Rep 8, 17608 (2018).

49. M. Orecchioni, Y. Ghosheh, A. B. Pramod, K. Ley, Macrophage Polarization: Different Gene Signatures in M1(LPS+) vs. Classically and M2(LPS–) vs. Alternatively Activated Macrophages. Front. Immunol. 10, 1084 (2019).

50. M. Papatriantafyllou, SOCS2 and SOCS3 in macrophage polarization. Nat Rev Immunol 13, 7–7 (2013).

51. M. F. Fontana, A. Baccarella, N. Pancholi, M. A. Pufall, D. R. Herbert, C. C. Kim, JUNB Is a Key Transcriptional Modulator of Macrophage Activation. J.I. 194, 177–186 (2015).

52. X. Tang, D. L. Marciano, S. E. Leeman, S. Amar, LPS induces the interaction of a transcription factor, LPS-induced TNF-α factor, and STAT6(B) with effects on multiple cytokines. Proc. Natl. Acad. Sci. U.S.A. 102, 5132–5137 (2005).

53. D. S. Hong, R. Kurzrock, J. G. Supko, X. He, A. Naing, J. Wheler, D. Lawrence, J. P. Eder, C. J. Meyer, D. A. Ferguson, J. Mier, M. Konopleva, S. Konoplev, M. Andreeff, D. Kufe, H. Lazarus, G. I. Shapiro, B. J. Dezube, A Phase I First-in-Human Trial of Bardoxolone Methyl in Patients with Advanced Solid Tumors and Lymphomas. Clin Cancer Res 18, 3396–3406 (2012).

54. R. Borella, L. Forti, L. Gibellini, A. De Gaetano, S. De Biasi, M. Nasi, A. Cossarizza, M. Pinti, Synthesis and Anticancer Activity of CDDO and CDDO-Me, Two Derivatives of Natural Triterpenoids. Molecules 24, 4097 (2019).

55. T. A. Stadheim, N. Suh, N. Ganju, M. B. Sporn, A. Eastman, The Novel Triterpenoid 2-Cyano-3,12-dioxooleana-1,9-dien-28-oic acid (CDDO) Potently Enhances Apoptosis Induced by Tumor Necrosis Factor in Human Leukemia Cells. Journal of Biological Chemistry 277, 16448–16455 (2002).

56. Y. Ito, P. Pandey, M. B. Sporn, R. Datta, S. Kharbanda, D. Kufe, The Novel Triterpenoid CDDO Induces Apoptosis and Differentiation of Human Osteosarcoma Cells by a Caspase-8 Dependent Mechanism. Mol Pharmacol 59, 1094–1099 (2001).

57. Q. Zhu, X. Wu, Y. Wu, X. Wang, Interaction between Treg cells and tumor-associated macrophages in the tumor microenvironment of epithelial ovarian cancer. Oncology Reports 36, 3472–3478 (2016).

58. X. Wang, M. Lang, T. Zhao, X. Feng, C. Zheng, C. Huang, J. Hao, J. Dong, L. Luo, X. Li, C. Lan, W. Yu, M. Yu, S. Yang, H. Ren, Cancer-FOXP3 directly activated CCL5 to recruit FOXP3+Treg cells in pancreatic ductal adenocarcinoma. Oncogene 36, 3048–3058 (2017).

59. M. Schenk, M. Fabri, S. R. Krutzik, D. J. Lee, D. M. Vu, P. A. Sieling, D. Montoya, P. T. Liu, R. L. Modlin, Interleukin-1β triggers the differentiation of macrophages with enhanced capacity to present mycobacterial antigen to T cells. Immunology 141, 174–180 (2014).

60. M. K. Yadav, Y. Inoue, A. Nakane-Otani, Y. Tsunakawa, H. Jeon, O. Samir, A. Teramoto, K. Kulathunga, M. Kusakabe, M. Nakamura, T. Kudo, S. Takahashi, M. Hamada, Transcription factor MafB is a marker of tumor-associated macrophages in both mouse and humans. Biochemical and Biophysical Research Communications 521, 590–595 (2020).

61. Y. Wang, C. Luan, G. Zhang, C. Sun, The transcription factor cMaf is targeted by mTOR, and regulates the inflammatory response via the TLR4 signaling pathway. Int J Mol Med (2018), doi:10.3892/ijmm.2018.3510.

