## Supplementary Material for "CDDO-Methyl Ester Inhibits BRAF Inhibitor Resistance and Remodels the Myeloid Compartment in BRAF-mutant Melanoma"

**SUPPPLEMENTARY DATA**

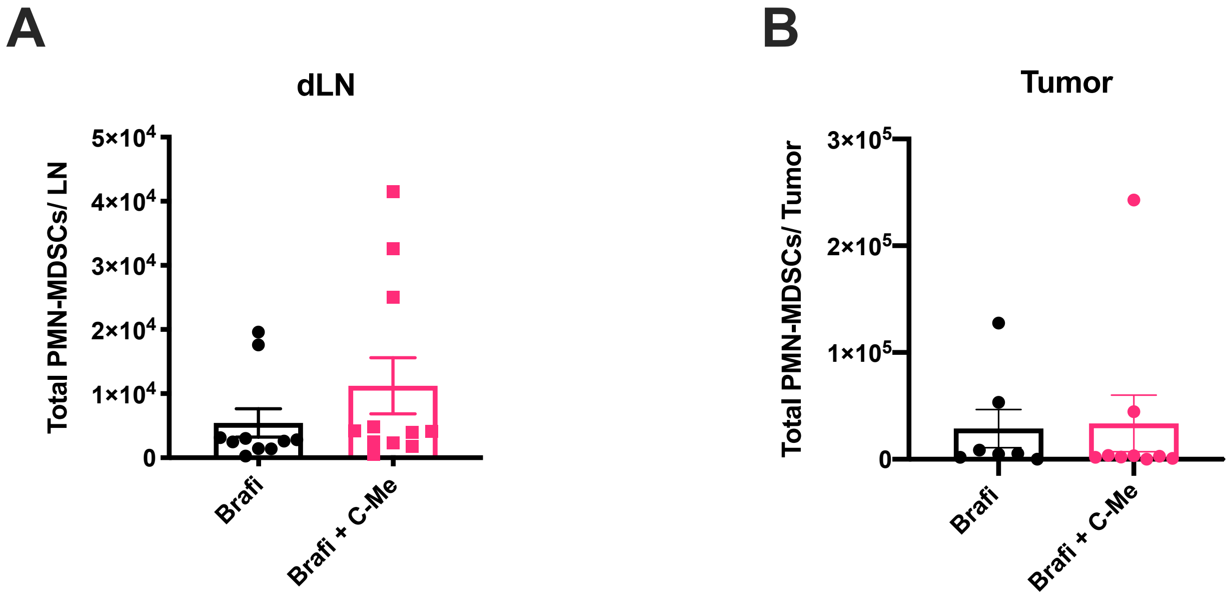


**Supplemental Figure 1: Total PMN-MDSCs in tumor draining lymph nodes and tumors.** (A) Draining lymph nodes (dLN) and (B) tumors from mono- (BRAFi) or dual-therapy (BRAFi + C-Me, 50 mg/kg) treated mice were harvested and analyzed by multicolor flow cytometry for analysis of the PMN-MDSC populations ($CD11b^+$, $Ly6C^-$, $Ly6G^+$). (A) Data are representative of 3 independent experiments involving 5-6 mice per group. (B) Data are representative of 3 independent experiments using 5-9 mice per group.

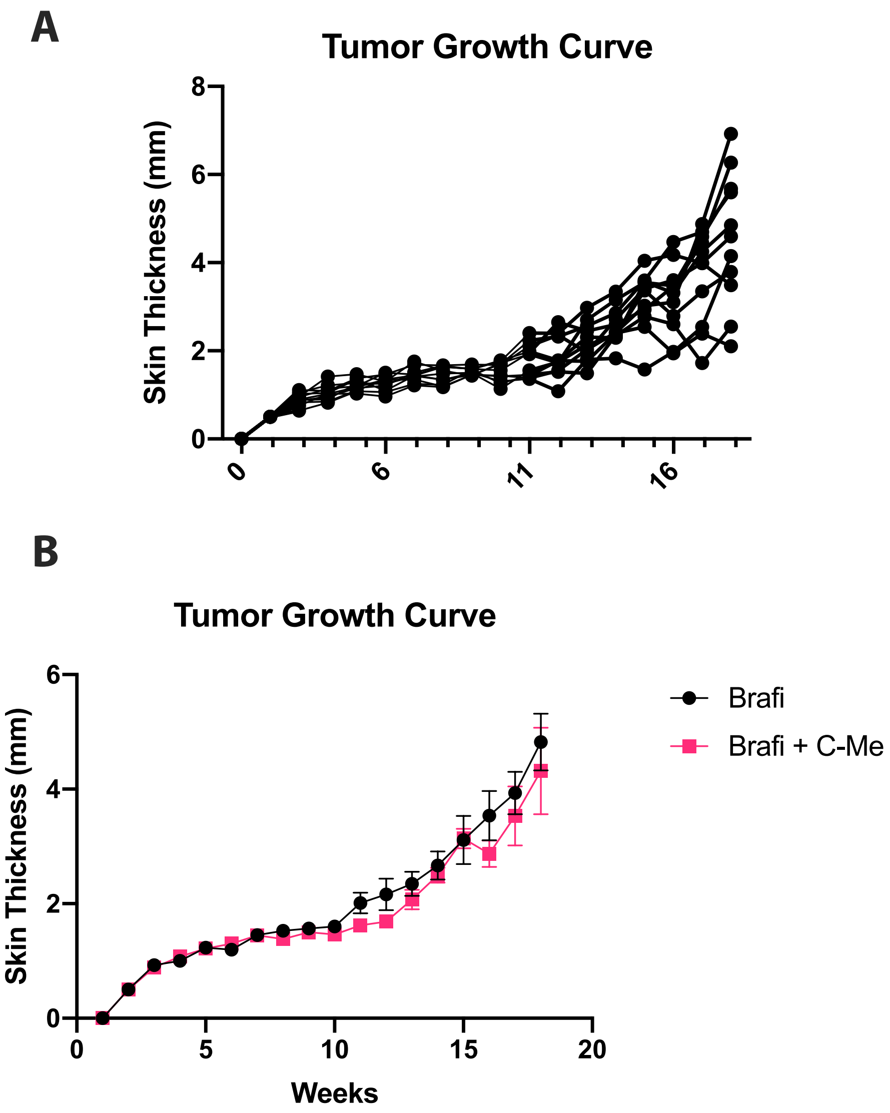


**Supplemental Figure 2: Tumor skin thickness measurements.** Tumors were measured for skin thickness weekly starting at the initiation of BRAFi treatment (see Figure 1) until the end of the experiment. (A) Individual mouse tumor thickness per week. (B) Average skin thickness per treatment per week.

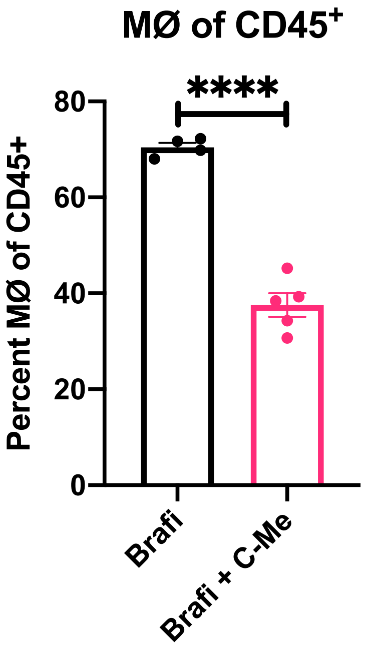


**Supplemental Figure 3: Total numbers of macrophage (MØ) in tumors.** Tumors from mice that received BRAFi or BRAFi/C-Me therapy (See Figure 1) were harvested and analyzed by multicolor flow cytometry for examination of macrophages (F4/80$^+$/CD45). Data shown are post-elimination of doublets and dead cells. Data were pooled from 3 independent experiments involving 4-5 mice per group. Statistical significance was calculated by two-tailed t-test. ****$p<0.0001$

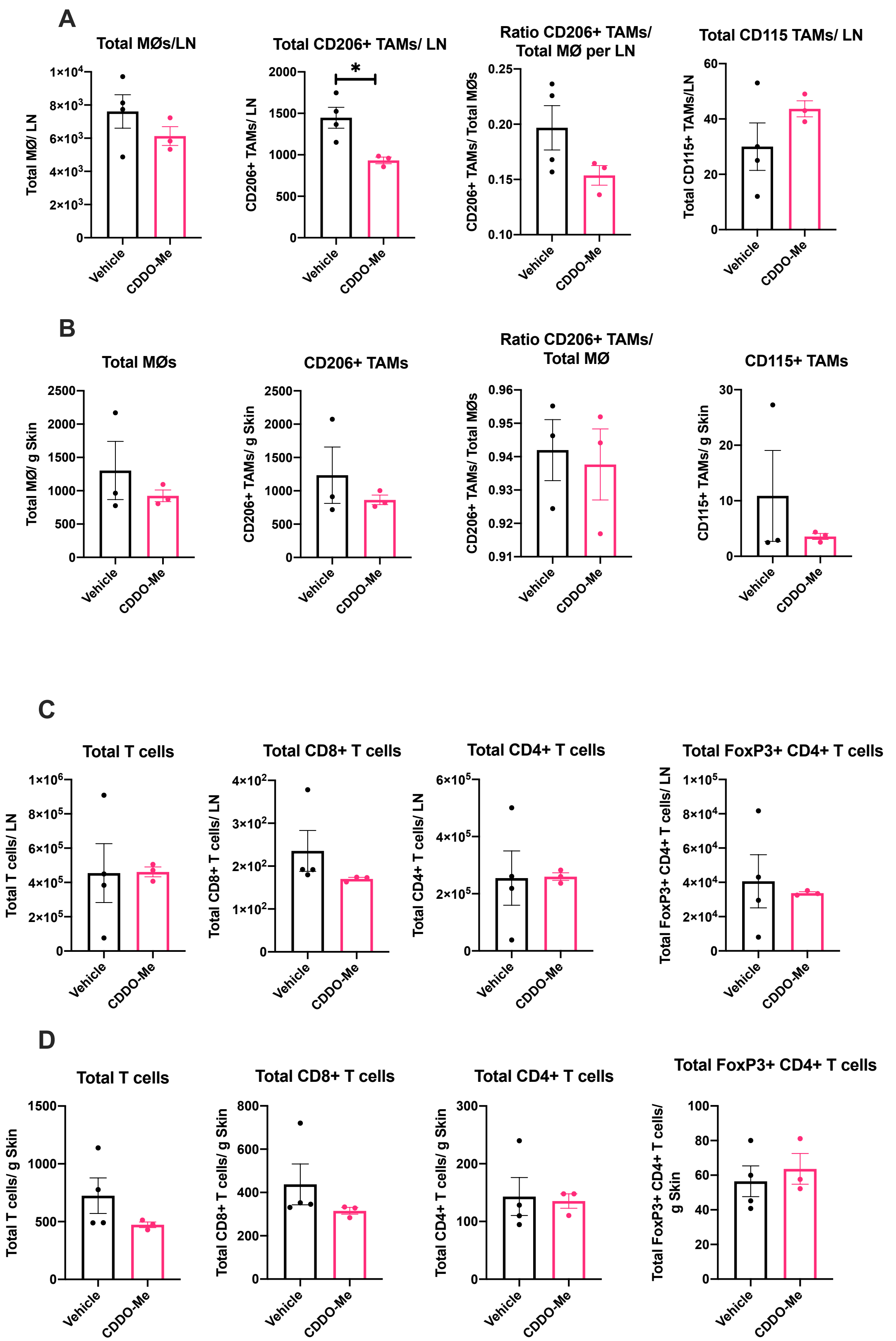


**Supplemental Figure 4 : C-Me monotherapy treatment in Braf/Pten mice post-4-HT tumor induction.** (A and C) dLN or (B and D) skin nevi from mice treated with vehicle or C-Me (50mg

C-Me/kg) for 26 days post-tumor induction. Mice were harvested and analyzed by multicolor flow cytometry for examination of (A-B) MØ and (C-D) T cell populations. Data shown are post-elimination of doublets and dead cells. Data were pooled from 3 independent experiments using 3-4 mice per group. Statistical significance was calculated by two-tailed t-test. *$p<0.05$

**Supplementary Table 1: All differentially expressed genes per cluster.**

https://docs.google.com/spreadsheets/d/e/2PACX-1vQNp1-eWGdy-B1vISSxDqrlepGC3l8E7q6fhZ_gmH8kCIjkdg-qFQUJsJCeqOYa9Czx4wiycwIxrm8F/pub?gid=1613746944&single=true&output=csv

**Supplementary Table 2: All differentially expressed genes per treatment.**

https://docs.google.com/spreadsheets/d/e/2PACX-1vSbwab22kthwNLDs_RpeBfDtdbicnGYJy2o06IhALVsRJp6fYkQjKv_7zIJIbN3cdn4yA17wxgJMNZq/pub?gid=94613830&single=true&output=csv
